# Functional redundancy in Oxa1-family insertases supports robust membrane insertion of influenza A virus M2 viroporin

**DOI:** 10.1101/2025.06.05.658076

**Authors:** Christian Diwo, Marta Alenquer, Shu Zhou, Paul Digard, Colin Adrain, Maria João Amorim

## Abstract

The biogenesis of thousands of highly diverse membrane proteins in humans is facilitated by an array of ER-resident membrane protein translocases. While some membrane proteins have a strict requirement for a specific insertion machinery, membrane proteins with short translocated domains may be able to access multiple pathways. Here, we quantify the functional importance of redundancy in membrane protein translocation during influenza A virus (IAV) infection by examining the biogenesis of the viroporin M2. Given the wide host and cellular tropism of IAV, the virus likely evolved mechanisms to leverage host translocation pathways efficiently. We demonstrate that although M2 utilizes the ER membrane protein complex (EMC), driven by signals encoded in its transmembrane and C-terminal domains, M2 maintains an approximately 50% membrane insertion rate in the absence of the EMC. This influences viral cell-to-cell transmission across different IAV strains, with a greater impact on those expressing lower levels of M2. We identify alternative translocation of M2 via Oxa1-family translocons independent of canonical targeting chaperones. These findings reveal how the exploitation of multiple redundant pathways can ensure robust IAV infection.

**SIGNIFICANCE STATEMENT:** IAV must rapidly replicate in diverse mammalian hosts, which requires efficient integration of viral proteins into host cell membranes. This study uncovers how the viral proton channel M2 utilizes multiple redundant protein insertion pathways, accessing EMC and alternative Oxa1-family translocases. Revealing these redundant strategies clarifies how cells triage membrane proteins, offering insights into both viral adaptation and host cell robustness.

## INTRODUCTION

Influenza A virus (IAV) is a major respiratory pathogen responsible for seasonal epidemics and occasional pandemics [1]. Rapid replication determines the success for transmission which requires efficient takeover of the host protein synthesis machinery [2–5]. The synthesis of membrane proteins presents a particular challenge due to the complexity of ER-targeting and insertion pathways [6] and may become rate-limiting for virion production.

Membrane protein biogenesis is shaped by biophysical properties—including transmembrane domain (TMD) hydrophobicity, flanking domain charges, secondary structure, and length—factors that determine pathway selection among ER-resident insertases [7–15]. Among the identified insertion sites are the Sec61 translocon, which accommodates the translocation of large hydrophilic domains, and Oxa1-family insertases such as the EMC, GET, and GEL complexes, which facilitate insertion of proteins with short terminal domains through a thinned lipid bilayer [6]. Despite various biophysical properties potentially triaging nascent membrane proteins to specific translocation sites, overlapping client pools have been identified [7,11,12,14]. The functional significance of this redundancy in potentially promoting robust membrane protein biogenesis remains unresolved.

IAV encodes three membrane proteins—hemagglutinin (HA), neuraminidase (NA), and matrix protein 2 (M2)—all expressed at the infected cell surface and incorporated into the viral envelope. While HA and NA are well-defined Sec61 substrates due to their large extracellular domains, the biogenesis of M2 remains poorly understood [16]. M2 is a 97-amino-acid, type III membrane protein [17,18] that forms a pH-sensitive tetrameric proton channel [19], with essential function during viral entry [20], and supporting roles during assembly [21–23], and budding [24]. M2 is the translation product of a spliced transcript from IAV genomic segment 7, and although virions require only small amounts of M2, it is often expressed at disproportionately high intracellular levels [25–27].

M2’s short C-terminal length and conserved hydrophilic residues in its transmembrane domain likely pose challenges for targeting and insertion into the ER membrane [28–31]. Due to its sequence features and its proven potential as a drug target [32], we investigated M2 biogenesis. We find that M2 utilizes EMC through signals encoded in its TMD and C-terminus. In the absence of the EMC, IAV virion production is delayed, depending on M2 expression level, which translates into a significant defect in cell-to-cell transmission. However, EMC-deficient cells retain substantial insertion efficiency, suggesting access to redundant insertion pathways. Our data reveal that alternative Oxa1-family translocons can compensate for EMC loss. This compensatory mechanism buffers viral replication and demonstrates how robustness in membrane protein biogenesis can arise from parallel pathways.

## RESULTS

### IAV M2 biogenesis is partially EMC dependent

Glycosylation is a prevalent post-translational modification, linking glycan chains to proteins entering the secretory pathway, which is widely used to assess membrane protein insertion into the ER and trafficking through the secretory pathway [33,34]. M2 needs to be inserted into the ER and trafficked to the plasma membrane to be incorporated into budding virions [35]. We track M2 membrane insertion via an N-terminally engineered glycosylation site through a single amino acid substitution (Leu3Asp) to obtain the reporter construct M2.Y.F (Y for a N-terminal glycosylation site and F for a C-terminal Flag-tag, Fig. 1A).

**Fig. 1.**
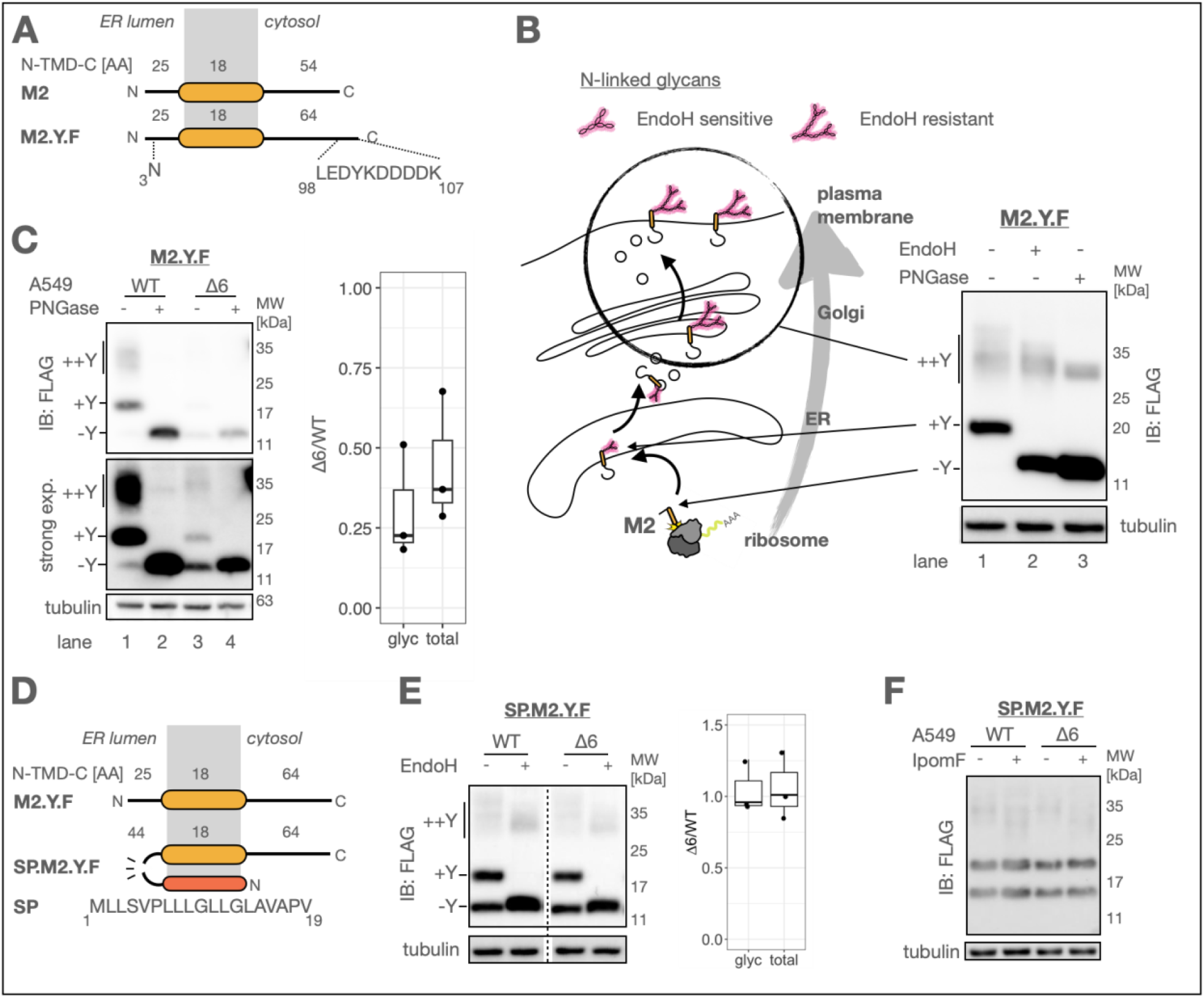
M2 is partially dependent on EMC for ER insertion. A) Scheme of M2 modifications, Leu3Asp and C-terminal Flag-tag to obtain glycosylation competent M2.Y.F B) Scheme and analysis of M2.Y.F trafficking along the secretory route and its resulting glycan modifications. A549 cells were transfected with M2.Y.F, and cell lysates were either mock-treated or deglycosylated with PNGase F or Endo H before analyzing by western blot (N=2). C) A549 WT or Δ6 cells were transfected with M2.Y.F and incubated for 24h. The cell lysates were analyzed by western blot. The glycosylation efficiency (glyc, +Y/SUM(+Y,-Y) and the total protein levels (total) were quantified from western blots by densitometry (N=3). D) Scheme of modification of M2.Y.F N-terminal fusion of calreticulin signal peptide to obtain SP.M2.Y.F. E) A549 WT or Δ6 cells were transfected with SP.M2.Y.F, and lysates were mock-treated or treated with Endo H and analyzed by western blot. The glycosylation efficiency (glyc, +Y/SUM(+Y,-Y) and the total protein levels (total) were quantified from western blots by densitometry (N=3). F) A549 WT or Δ6 cells were transfected with SP.M2.Y.F and incubated with 50nM Ipomoeassin F (IpomF), or equivalent volume fraction of DMSO and lysates were analyzed by western blot (N=3).

Western blot analysis of A549 lung cancer cells transfected with M2.Y.F reveals two distinct populations at steady state (Fig. 1B lane 1). Differential glucosidase digestion with Endo H (only cleaves high mannose N-linked glycans) or PNGase F (cleaves all N-linked glycans) reveals these populations to be trafficking intermediates corresponding to ER membrane inserted M2 (glycosylated +Y at 19kDa, Fig. 1B), or M2 transiting the Golgi (diverse glycan modifications ++Y around 35kDa, Fig. 1B).

Previous research has proposed that most type III membrane proteins are inserted into the membrane via the EMC [12,13]. Therefore, to assess the impact of loss of the EMC on M2 membrane insertion, we generated A549 cells deficient in EMC6 (Δ6), a core subunit of the EMC whose loss ablates EMC function, and results in the loss of other EMC subunits [36] (Fig. S1). Δ6 cells glycosylate HLA-A (N-terminal FLAG-tag, Sec61 client), and SEC61B (C-terminal HA-tag, GET client), but do not glycosylate a known EMC client SQS (C-terminal HA-tag, Fig. S1). Reconstituting EMC6 in the Δ6z cells (+6) rescues SQS glycosylation (Fig. S1). Notably, M2.Y.F transfected into WT and Δ6 cells revealed a decrease in the overall levels (Fig. 1C lane 2 and 4) and a shift in the proportions of an unglycosylated band (-Y) over +Y at steady state (glycosylation efficiency +Y/SUM(+Y,-Y), Fig. 1C lane 1 and 3). Overall protein levels were reduced to ~30% Δ6/WT and glycosylation efficiency was reduced to ~25% Δ6/WT (Fig. 1C). The increase in −Y in the absence of EMC implies a defect in M2 biogenesis. M2 may partially be unglycosylated due to its N-terminus remaining in the cytosol, either due to a defect in membrane insertion, failed topogenesis, or M2 perhaps erroneously insert into other organelles in the absence of EMC [37].

N-terminal signal peptide encoding proteins, like calreticulin, enter the ER through Sec61 [38]. We fused the signal peptide of calreticulin to M2.Y.F’s N-terminus (SP.M2.Y.F, Fig. 1D) to redirect M2 biogenesis to Sec61, thus testing if EMC may indirectly act on M2 biogenesis through stabilizing other biogenesis factors [39]. SP.M2.Y.F alleviated a defect in glycosylation efficiency and expression level in Δ6 compared to WT cells, arguing for EMC directly mediating M2 targeting or translocation (Fig. 1E). We tested if SP.M2.Y.F was indeed redirected to be inserted by Sec61 by inhibiting lateral gate opening with Ipomoeassin F (IpomF) [40]. IpomF inhibits PR8 HA expression (Fig. S2), however it did not impact the glycosylation efficiency or overall protein levels of SP.M2.Y.F in WT or Δ6 cells (Fig. 1D). M2’s N-terminus is partially unstructured [41], a feature that has been reported to hinder Sec61 translocation despite the presence of a functional signal peptide [9,42,43]. We hypothesize that a signal peptide influences SP.M2.Y.F’s targeting dynamics to the ER membrane, which results in EMC and Sec61 independent insertion through another translocon. We conclude that the EMC is likely to directly act early in M2.Y.F biogenesis, either by increasing targeting efficiency or by directly mediating ER membrane insertion.

### Loss of the EMC reduces M2’s plasma membrane accumulation rate and the onset of virion production

Next, we tested whether EMC-function also affects the fate of native M2 during IAV infection. Infection with the lab-adapted IAV strain A/Puerto Rico/8/34 (PR8) for 8 and 12 hours showed a sustained loss of M2 relative to virally-encoded nuclear protein (NP), normalized to a loading control (~25% Δ6/WT) (Fig. 2B). Although overall M2 levels were reduced—consistent with observations for other bona fide EMC clients [14,39,44]—residual M2 levels slightly increased with time in Δ6 cells until 12 hours post-infection (hpi) (Fig. 2B).

**Fig. 2.**
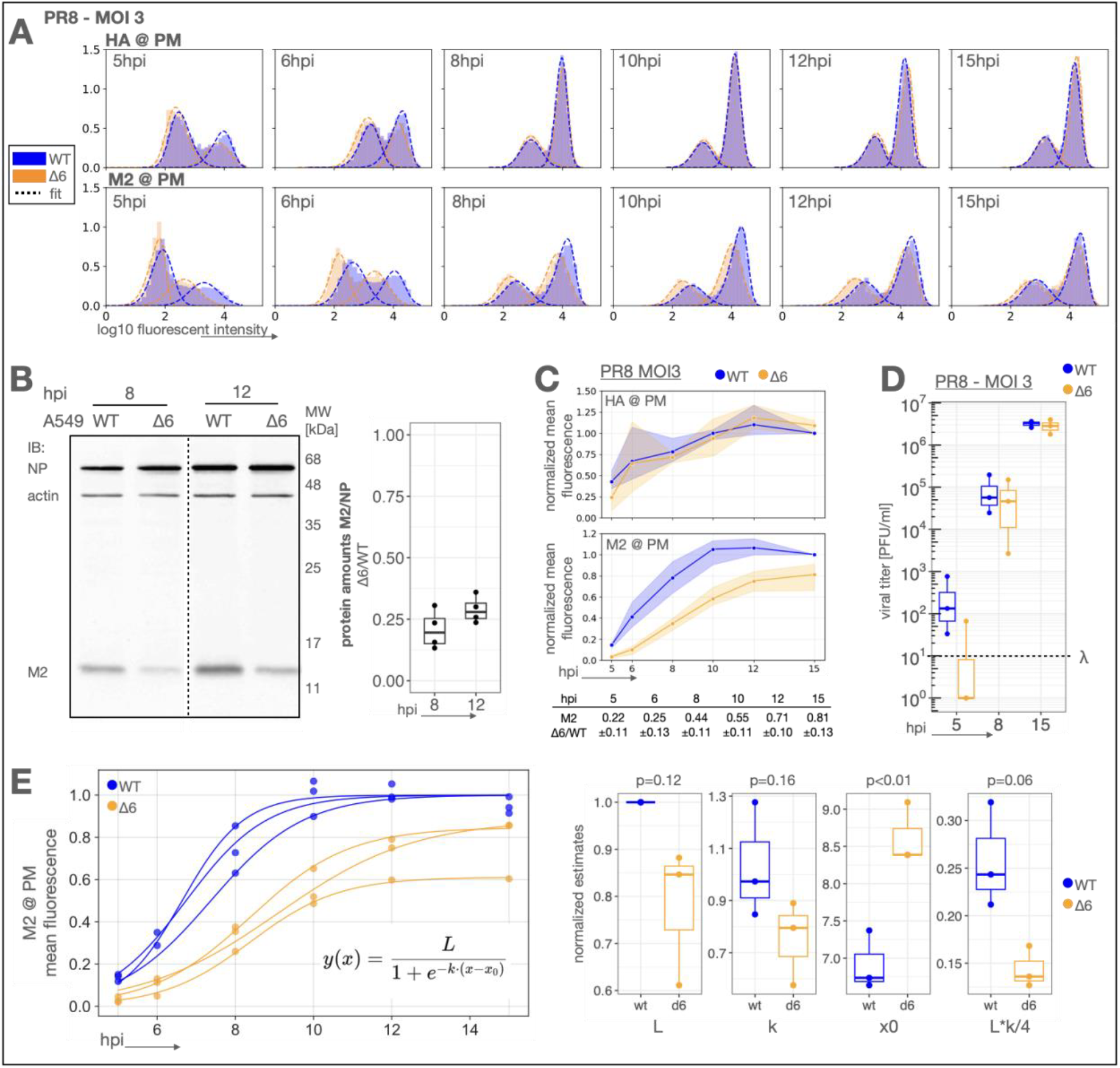
EMC ablation affects M2 trafficking dynamics in infection. A) A549 WT or Δ6 cells were infected with PR8 at MOI 3 for cell harvest at the indicated time points and subsequent surface expression of viral HA and M2 by flow cytometry. The mean of an HA or M2 positive population of single cells was accurately estimated by fitting a skew-normal distribution (N=3). B) A549 WT or Δ6 cells were infected with PR8 at MOI of 3, and cell lysates were harvested 8 or 12 hours post infection. The total protein levels were quantified from western blots by densitometry (N=4). C) Time series of the estimated means from A) for HA and M2 normalized to the value WT conditions at 15hpi for each protein in each replicate. D) Viral titre analyzed by plaque assay from supernatants of the 5h, 8h, and 15h time points (N=3). λ denotes the limit of detection in this assay. E) A logistic growth model was fitted to each repeat of time series, normalized to the estimated parameter L in WT conditions and fit again to the normalized data. The maximal slope corresponds to the maximal rate of accumulation, and is calculated from the estimated parameters L*k/4.

EMC functions upstream in M2 biogenesis, before M2 trafficking reactions to reach the plasma membrane. We quantified the effect of EMC loss on M2 in infection by immuno-detecting M2’s N-terminus at the plasma membrane in a flow cytometry assay over the course of infection from 5 to 15hpi (Fig. 2A). Interestingly, we observed a defect in M2 accumulation at the plasma membrane, which was most pronounced at 5hpi (22% Δ6/WT) but approached WT levels by 15hpi (81% Δ6/WT) (Fig. 2C). The virally encoded Sec61-dependent protein HA was not affected by loss of EMC function, congruent with a specific defect in M2 biogenesis (Fig. 2C). These results argue against a defect in viral entry or mRNA expression due to the loss of EMC.

Fitting the time series of M2 plasma membrane accumulation to a logistic growth model provides interpretable parameters, such as the maximal accumulation rate of M2 at the plasma membrane, which was reduced to 56% in the absence of EMC compared to WT conditions (Fig. 2E). Surprisingly, despite the pronounced defect in overall M2 protein levels and plasma membrane accumulation rate, viral titre at 8 and 15hpi were unaffected (Fig. 2D). However, the delay in M2 accumulation at the plasma membrane correlated with a delay in the onset of virion production as most Δ6-infected samples lacked detectable levels of infectious virus particles earlier, at 5hpi, whereas all WT samples produced virus at this time point (Fig. 2D). We concluded that the loss of EMC affects M2 levels in infection, reducing the rate of M2 accumulation at the plasma membrane. The reduced rate of M2 trafficking correlates with a delayed onset of virion production, however, M2 reached the plasma membrane in sufficient levels to support normal virion production by 8hpi in PR8 infection. These results consolidate that EMC is important for M2 biogenesis, yet also highlight a striking compensation for EMC loss by an unidentified alternative insertion pathway.

### Cell-to-cell transmission is impacted by M2 biogenesis defect

Rapid cell-to-cell transmission is crucial to quickly produce large quantities of infectious virus particles in an infected host [45]. Our results indicate that the onset of M2 expression shapes the dynamics of virion production, which may in turn affect cell-to-cell transmission. To test this, we resorted to MDCK cells as a model. MDCK cells strongly adhere to cell culture dishes and tolerate the moderate concentrations of trypsin in the media that are required to cleave PR8 HA for multicycle viral infection [46]. In a single-cycle infection at MOI 3, MDCK Δ6 cells infected with PR8 virus selectively lost most M2 expression without affecting other measured viral proteins (Fig. 3A), and we observed a modest growth defect at 6hpi (Fig. 3B). Both M2 expression and viral growth were rescued in +6 cells (Fig. 3B). M2 expression levels differ between viruses due to variation in M gene splicing (segment 7) and may be adapted to the host species [4,47]. We hypothesized that viruses expressing lower levels of M2 may show an increased EMC-dependent delay in virion production. We used a PR8 segment 7 splice site acceptor mutant D232Q (G719C+T721G) that retained normal M1 expression but downregulated M2 expression (Fig. S3). Indeed, a virus that expressed limiting levels of M2 showed an increased delay in virion production due to EMC loss (Fig. 3C), consolidating M2’s role in shaping virion production dynamics.

**Fig. 3.**
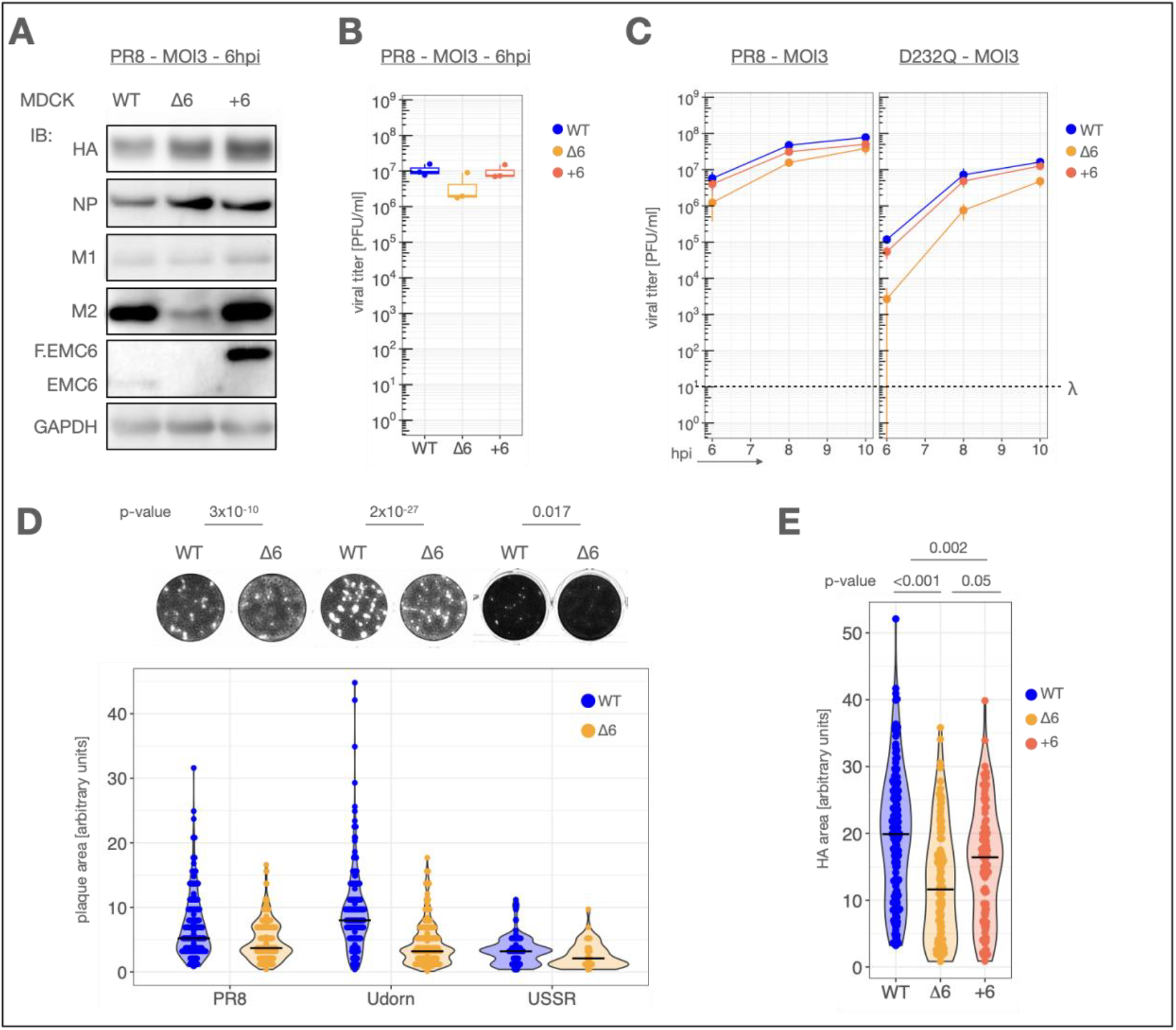
M2 biogenesis defect impacts cell to cell transmission in MDCK cells. A) MDCK WT, Δ6 or +6 cells infected with PR8 at MOI of 3 for 6 hours. Cell lysates are analyzed by western blot (N=2). B) Supernatants were titrated by plaque assay (N=3) C) MDCK WT, Δ6 or +6 cells infected with PR8 or D232Q segment 7 splice site mutant at MOI of 3 for indicated time points and supernatants were titrated by plaque assay (N=2). λ denotes the limit of detection in this assay. D) A confluent monolayer of MDCK WT or Δ6 cells was infected with 60 PFU of PR8 virus and incubated with viscous AVICEL overlay for 30h. Cells were fixed, stained, imaged and plaque area was quantified in imageJ (N=6). P-values were obtained by t-test.E) A549 WT, Δ6 and +6 cells were infected as in B), fixed and stained with PR8-HA antibody and imaged. Area was quantified in imageJ and p-values assessed by one-way ANOVA (N=2).

To assess cell-to-cell transmission, we infected monolayers of MDCK WT or Δ6 cells with a low titre of virus (60 PFU per well) and quantified plaque area at 30hpi (Fig. 3D). We tested H1N1 virus PR8, a lab-adapted H3N2 virus A/Udorn/307/1972 (Udorn), plus a clinical isolate A/USSR/90/77 (USSR77) [48]. All viruses were significantly affected by the loss of EMC in their cell-to-cell transmission, as determined by plaque size (Fig. 3D), which is in agreement with other studies limiting M2 at the infected cell surface [35,49]. We confirmed that the measurements of plaque area correlates with cell-to-cell transmission by assessing the area of cells expressing HA for PR8 infected cells (Fig. 3E). Udorn, a fast-growing, filamentous virus demonstrated a more significant defect in cell-to-cell transmission than PR8 in this assay. Udorn M2 has 10 amino acid substitutions compared to M2 of PR8 but retains EMC dependence (Fig. S4). These findings are relevant as filamentous virus morphology is prevalent in patient samples [50,51] and may aid cell-to-cell transmission [52,53]. In conclusion, our results show that alterations in the dynamics of virion production due to a defect in M2 biogenesis can significantly slow cell-to-cell transmission.

### M2 sequence features determine biogenesis route at the ER

Transmembrane domain hydrophobicity and C-terminal length have both been identified as potential signals for EMC dependence [7,12,13]. M2 (indicated with the red line) has a relatively hydrophilic transmembrane domain (ΔGapp −0.09 kcal/mol) and a cytosolic C-terminus that is very short (54 amino acids) compared to the human membrane proteome (Fig. 4A, M2 characteristics indicated by red dashed lines). Based on these characteristics, we searched the human proteome for a type III protein with a hydrophobic transmembrane domain and long C-terminus. SRPRB, a highly conserved membrane receptor for co-translational targeting, fit these criteria with a hydrophobic transmembrane (ΔGapp −1.8 kcal/mol) domain and a longer C-terminus (226 amino acids) (Fig. 4A, SRPRB characteristics indicated as purple dashed lines). SRPRB’s N-terminus was mutated to contain a glycosylation site and its biogenesis determined to be EMC independent, making it an ideal model to test sequence features of EMC dependence (Fig. 4B). Subjecting domain-swap chimeras between M2 and SRPRB to the same glycosylation assay described in Fig. 4B, we found M2’s C-terminus and transmembrane domain carried signals of EMC dependence, affecting translocation efficiency and protein levels in absence of EMC (Fig. 4B). We did not observe an effect of EMC loss on a construct carrying M2’s N-terminal domain (MSS.Y.F, Fig. S5).

**Fig. 4.**
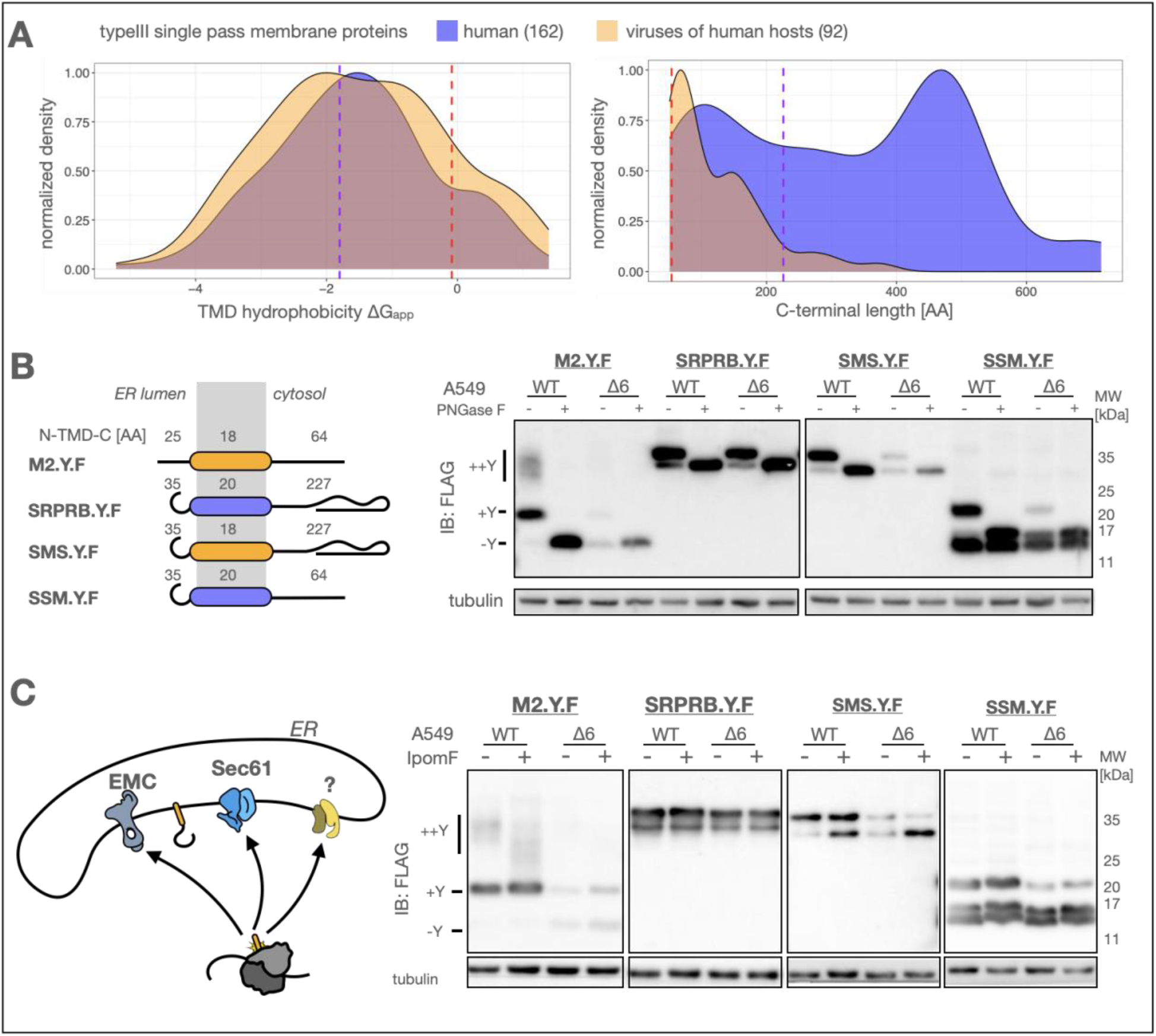
M2 sequence features determine biogenesis route at the ER. A) Single-pass type III proteins of the human proteome (162), or proteomes of viruses of human hosts (92) were analyzed for transmembrane domain hydrophobicity (ΔGapp [10]) and length of C-terminal domain. Red dashed lines indicate M2 characteristics, purple dashed lines indicate SRPRB characteristics. B) A549 WT or Δ6 cells were transfected with each of the indicated constructs, and cell lysates were mock-treated or treated with PNGase F and analyzed by western blot (N=3). C) A549 WT or Δ6 cells were transfected with M2.Y.F, SRPRB.Y.F, SMS.Y.F or SSM.Y.F and incubated with 50nM IpomF or equivalent volume fraction of DMSO for 24h, and lysates were analyzed by western blot (N=3).

Type III proteins may engage both EMC and Sec61, depending on the length of the co-translationally targeted nascent chain interacting with the ER membrane [11]. We tested if any of our type III proteins are Sec61 dependent by blocking Sec61 insertion with IpomF. Interestingly, only SMS.Y.F (SRPRB carrying M2’s transmembrane domain) showed sensitivity to IpomF in WT and EMC ablated conditions, without inhibiting all insertion (Fig. 4C). These results show that type III protein biogenesis pathway is directed by diverse sequence features and that access to multiple pathways, or strict selectivity, is a feature likely be tuned by sequence context.

### M2 C-terminal length influences EMC dependence and redundant translocation pathway

Transmembrane domain features have been tested extensively to influence membrane insertion at the ER. C-terminal length has only recently been suggested to also contribute to pathway selection in membrane protein biogenesis. To test this in our system, we constructed M2 mutants where we concatenated the same C-terminal 37 amino acids (residue 60 to 97, conserving the same nucleotide sequence) to achieve a longer C-terminus that conserves potential short linear motifs (Fig. 5A). We observed that the C-terminal extension of M2 progressively gained glycosylation efficiency (Fig. 5A). The gain in EMC independence due to C-terminal extension might be due to the elongation of the mRNA, however, the EMC dependence of SMS.Y.F, which is also expressed from an elongated transcript would argue against mRNA length playing a role. As was the case for SMS.Y.F, the M2 C-terminal extension mutants become partly dependent on Sec61 processing, albeit to a very low degree, judged by the appearance of an unglycosylated band under IpomF treatment (Fig. 5B). These results argue that the extension of translation time after signal emergence influences EMC dependence and the selection of a translocon, which is in line with our earlier observation that SP.M2.Y.F becomes EMC independent, effectively extending the sequence length after signal emergence.

**Fig. 5.**
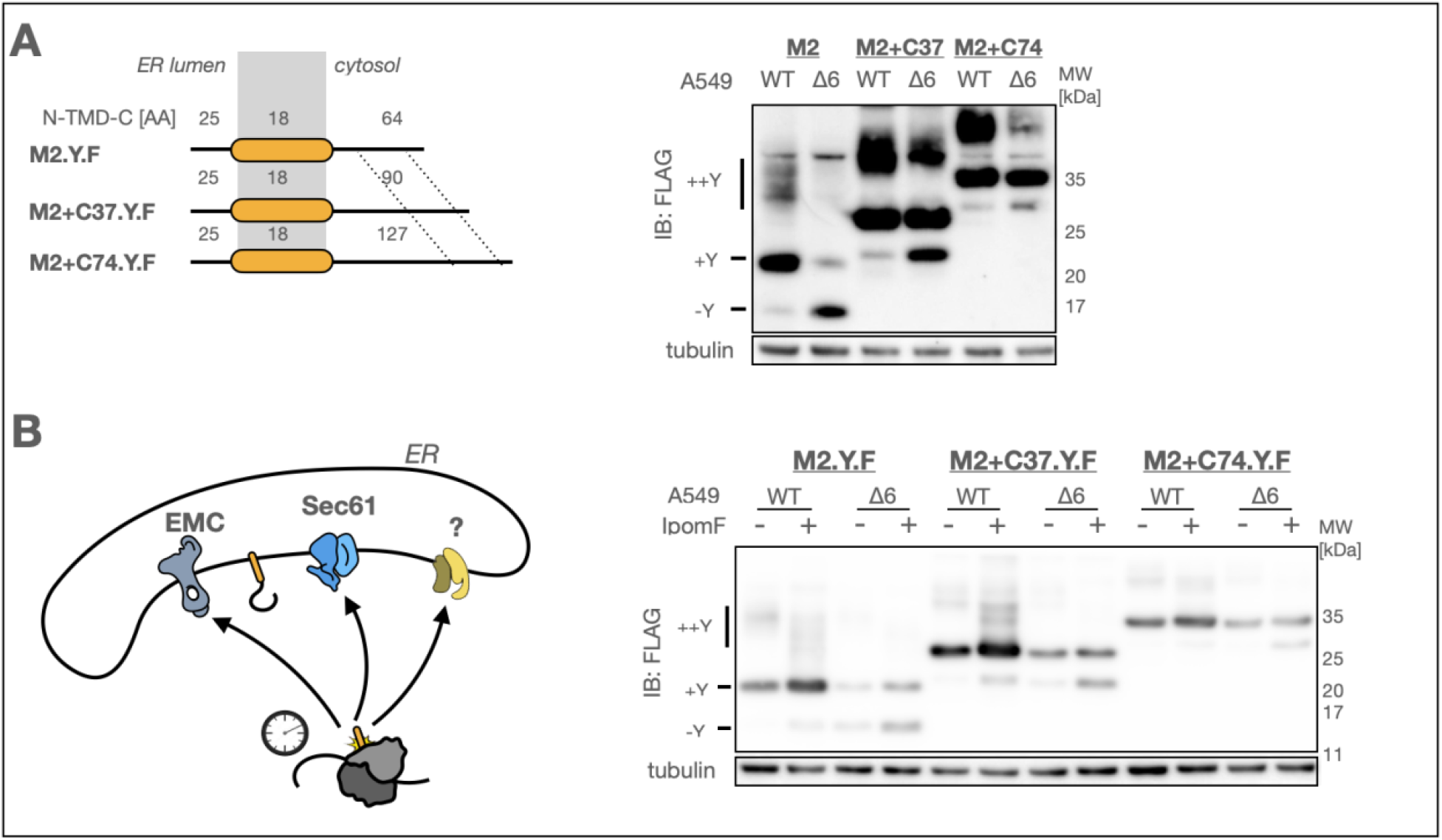
M2 C-terminal length determines the translocation pathway. A) Scheme of C-terminal extension mutants, where the last 37 amino acids of M2’s C-terminal were concatenated. A549 WT or Δ6 cells were transfected with each of the constructs for 24h and analyzed by western blot (N=2). B) A549 WT or Δ6 cells were transfected with M2 C-terminal extension mutants and incubated with 50nM IpomF or equivalent volume fraction of DMSO and lysates were analyzed by western blot (N=3).

### Oxa1-family members act as redundant M2 translocons

Our results indicate a relatively high rate of insertion in absence of EMC, which is likely mediated by an alternative translocon. As Sec61 does not act as redundant insertase we tested alternative Oxa1-family members, GET and GEL. To do so we targeted subunits of the Oxa1-family translocons by CRISPR KO in 293T cells (Fig. 6A). We confirmed population-level target depletion via western blot 72h after guide transfection (Fig. 6B).

**Fig. 6.**
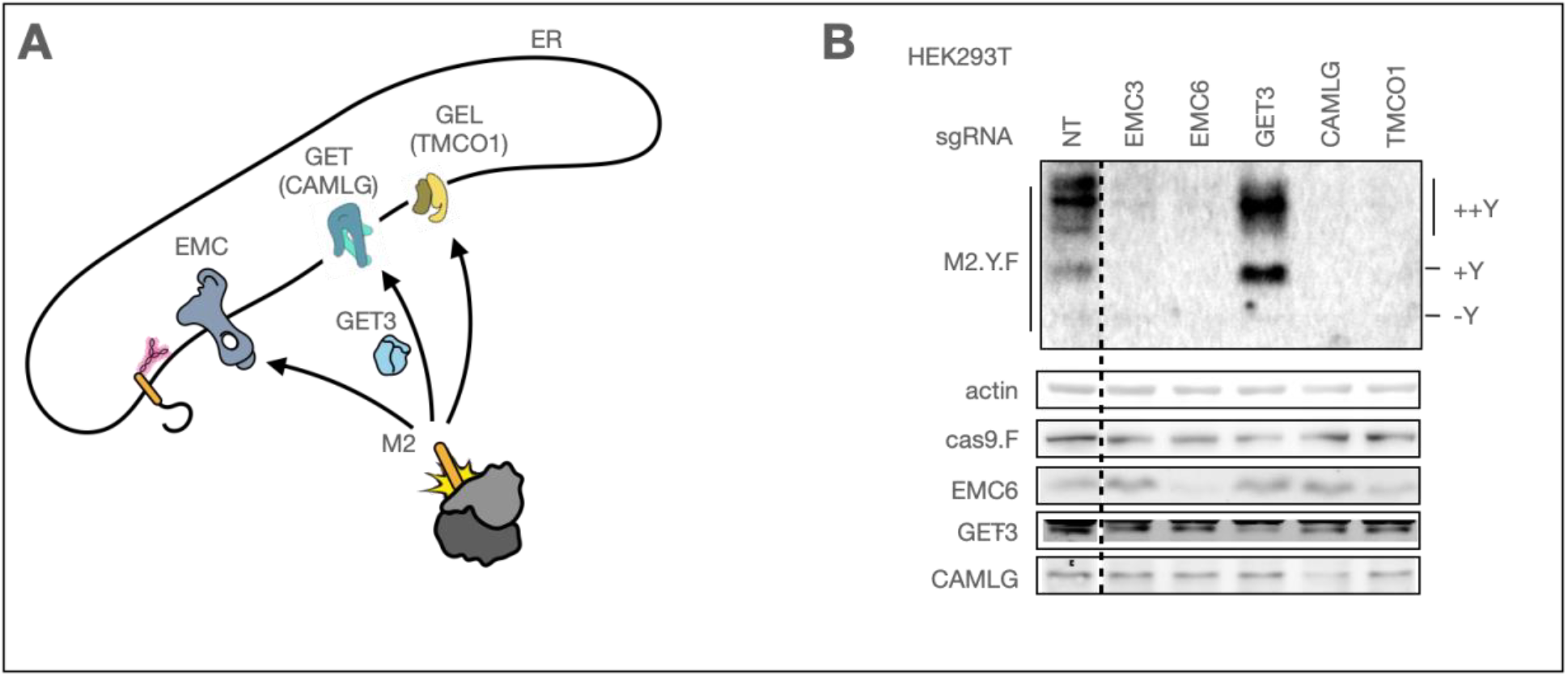
Oxa1-family members act as redundant M2 insertion sites. A) Scheme of targeted Oxa1-family insertase subunits. B) HEK293T cells were transfected with cas9 and subsequently with in vitro transcribed guides against a specific component of the membrane protein targeting or translocation machinery. KO cells were transfected with M2.Y.F, and the cell lysates analyzed by western blotting (N=2).

Surprisingly, in addition to our established dependency on the EMC (EMC3 and its obligatory cofactor EMC6), M2.Y.F levels were affected by several Oxa1-family members, the GET complex (CAMLG [15]) and GEL (TMCO1 [54]) components, but not the GET pathway-specific targeting factor GET3 (reviewed in Hegde and Keenan, 2024 [6]). To date, no targeting factor has been described to cater all Oxa1-family insertases [6]. Taken together, our data suggests that M2.Y.F can utilize multiple Oxa1-family members for its membrane insertion, through a targeting mechanism which does not depend on GET3.

## DISCUSSION

Successful IAV transmission depends on rapid virion production in a host, before immune reactions can limit infectious virus particle shedding [2,3,5]. Viral membrane proteins may become rate limiting virion components due to the complexity of their synthesis. IAV expresses three membrane proteins, NA and HA, which strictly require the Sec61 translocon and M2, whose biogenesis pathway was previously unknown. Our work illuminates the molecular details of how the critical virion component M2 is inserted into the ER membrane. Against the prevailing understanding of client-specific membrane protein biogenesis pathways, we reveal M2’s capacity to access several redundant translocons of the Oxa1-family. This redundancy allows efficient viral replication in absence of the abundant ER translocon EMC, demonstrating for the first time how redundancy in membrane protein biogenesis pathways promotes robustness.

Our findings show that even very low levels of M2 at the plasma membrane are enough to support IAV replication. Notably, although M2 levels drop to just 25% in EMC-deficient cells, PR8 virus titers are only affected at the earliest time points. Mutations in segment 7 that reduced M2 expression exacerbated the early defect in virion production, but also these viruses reached wild type-level titre later in infection, indicating that M2 is only rate limiting early in infection. However, when M2 biogenesis is impaired by removing the EMC, we observe a defect in cell-to-cell transmission across multiple IAV strains, highlighting the role of M2 in shaping infection dynamics. These observations have implications for M2 as an antiviral target, as splice site strength can rapidly evolve [55], which may render therapies aimed at limiting M2 expression ineffective. This may also explain how previous screens for IAV host-factors have missed identifying EMC [56–60], despite EMC being important for diverse flaviviruses [61,62]. Importantly, human immunodeficiency virus (HIV) ion channel Vpu has recently been investigated for its biogenesis requirements and found to be EMC-dependent [13]. In our analysis of proteomes of viral genera containing human-infecting species we identified 93 diverse type III proteins expressed in viruses of 22 different genera (Table S1), which may render these viruses EMC-dependent.

While membrane proteins with large translocated domains have a strict requirement for Sec61 processing, membrane proteins with short translocated domains may be more flexible to either use Sec61 or Oxa1-family insertases. Building on previous observations [12,13], we identified the low hydrophobicity and short C-terminal length of M2 as influencing sensitivity to EMC ablation and Sec61 lateral gate inhibition. Our results suggest that the targeting modality of M2 to the ER may dictate the selection of a translocon. Extension of translation time after signal emergence either by fusion of an N-terminal signal peptide, or C-terminal extension decreased sensitivity to EMC ablation. Furthermore, signal anchor hydrophobicity may influence the propensity of SRP targeting [63] or handling between cytosolic chaperones [64]. Further research is needed to understand how targeting mechanisms in conjunction with nascent membrane protein features expand or restrict the available repertoire of parallel membrane protein translocation pathways.

### LIMITATIONS OF THIS STUDY

Our results show that M2 biogenesis is affected by the loss of EMC, GET or GEL subunits, however we do not confirm a direct interaction between M2 and these translocons, which is expected to be very transient.

The discrepancy between total M2 levels measured by western blot and M2 levels at plasma membrane may be due to cells lost during washing and centrifugation steps in our flow cytometry preparations.

## MATERIALS AND METHODS

### Cell lines

A549 (ATCC CCL-185), HEK293T (ATCC CRL-3216) and Madin-Darby Canine Kidney (MDCK.1, ATCC CRL-2935), were cultured in Dulbecco’s Modified Eagle Medium (DMEM, Thermo Fisher 21969-035) supplemented with 10% fetal bovine serum (FBS, Thermo Fisher 10500064), 2 mM L-glutamine (Thermo Fisher, 25030024), and 1% (v/v) penicillin-streptomycin (Biowest L0022-100) in a humidified incubator at 37°C and 5% CO_2_.

### Antibodies

anti-Flag-tag (Sigma-Aldrich F1804-200UG), anti-HA-tag, anti-M2 14C2 (Abcam ab5416), anti-PR8-HA hybridoma (gift from Jonathan Yewdell), anti-NP (Abcam ab128193), anti-M1 (Abcam ab20910), anti-EMC6 (Abcam ab84902), anti-EMC5 (Abcam ab122202), anti-tubulin YL1-2 hybridoma (homemade), anti-ACTB (Sigma-Aldrich A5441), anti-GET3 (Abnova H00000439 M03), anti-CAMLG (Cell Signalling Technology 16713913S), anti-GAPDH (Sicgen, AB0049)

### Reagents

Avicel (Merck Supelco 11365-1KG), Formaldehyde (Acros 10231622), JetPrime (Polyplus 101000046), Fugene HD (Promega PROME2313), ECL Select (GE Healthcare RPN2235), BSA (Sigma-Aldrich A9418-100G), PNGase F (New England Biolabs 174P0704S), Endo H (New England Biolabs P0702), nitrocellulose membrane (GE Healthcare 10600003), culture plates (Corning 734-1597), poly-D-lysine (Sigma-Aldrich P1024-100MG), Bradford assay (Sigma-Aldrich B6916), Phusion (NEB 174M0530L), Pfu Ultra (Agilent Technologies 600387)

### Cell line construction

Polyclonal knockouts of EMC6 were obtained by transfecting A549 cells with 3 plasmids pX330 encoding the respective on-target sgRNA, or non-targeting sgRNA together with a plasmid for puromycin selection using Fugene HD transfection reagent. 24 hours post transfection, cells were selected with 3ug/ml puromycin for 72h or until all control cells not receiving selection plasmid died. For A549 WT and EMC6 KO cells the cells were subjected to another round of transfection and selection to obtain a polyclonal population. To obtain oligoclonal MDCK WT and EMC6 KO cells, the cells were sorted into 96-well plates, and clones were selected following verification of protein depletion by western blotting. 10 WT and EMC6 KO clones were combined to obtain an oligoclonal population. Rescue cell lines were obtained by lentiviral transduction with a pLEX either containing a FLAG-tag open reading frame, or N-terminally tagged FLAG.EMC6 and selected in hygromycin for 72h or until all control cells not receiving lentivirus died.

### Viruses

A/Puerto Rico/8/34 (PR8) virus were obtained via reverse genetics plasmid system (de Wit et al., 2007). 8 plasmids were transfected into 293T cells, and supernatants were used to expand in MDCK cells. The final viral stock solutions were obtained by using the supernatants from infected MDCK cells to infect embryonated chicken eggs. PR8 segment 7 D232Q was produced in the same way using the mutated segment. A/Udorn/307/1972 and A/USSR/90/1977 were a gift from Prof. Paul Digard, Roslin Institute, UK.

### Plasmids

Primers used for each construct are listed in Table S2. Plasmids for SEC61B.H and SQS.H were a gift from John Christianson’s lab [14]. HLA-A.F was amplified from human cDNA and cloned into pcDNA3 with a C-terminal Flag-tag and an AVI-tag following the endogenous signal peptide. M2 was amplified from cDNA generated from PR8 infected A549 cells and cloned into pcDNA3 with a C-terminal Flag-tag using KpnI and XhoI restriction sites. The Leu3Asp mutation generating the glycosylation site was introduced by quick change mutagenesis yielding M2.Y.F. SRPRB was amplified from mouse cDNA and cloned into pcDNA3 with a C-terminal Flag-tag using KpnI and XhoI restriction sites. The glycosylation site at SRPRB’s N-terminus was introduced so that the protein sequence starts with amino acids MNGT. Domain chimeras between SRPRB and M2, MSS.Y.F, SMS.Y.F, and SSM.Y.F were achieved by overlap-PCR and cloned into pcDNA3. SP.M2.Y.F was cloned by adding the calreticulin signal peptide to M2.Y.F by overlap-PCR. PR8-HA was amplified from the PR8 reverse genetics system and cloned into pLEX. cas9-BLAST was ordered from Addgene (52962).

### Transfection, western blots and deglycosylation

A549 WT or Δ6 cells were seeded in 24 well plates (6×10^4 cells/well) and transfected 16h later with plasmid (each 150ng/well) with Jetprime reagent and incubated for 20h. Cells were lysed in 70ul TX100 lysis buffer (50mM Tris-HCl pH 7.4, 150mM NaCl, 1% (v/v) Triton X-100, 5mM EDTA and protease inhibitors) on ice, scraped and transferred to 1.5ml test tubes and spun at 1000xg and 4°C. The supernatant was transferred to new tubes, mixed with 6x LDS sample buffer (550 mM Tris-HCl pH 7.4, 200 mM LDS, 0.13 mM EDTA, 250 mM DTT, 15% v/v glycerol, 0.025% SERVA Blue G250 and 0.025% Phenol Red) and to a 95°C hot plate for 3min. Samples were stored at −20°C until analysis by western blot.

If needed a small amount of sample was deglycosylated with NEB PNGase F (3%v/v per reaction) or Endo H (3%v/v reaction) as per manufacturer’s description. 10% Acrylamide gels containing SDS and sucrose were loaded with samples and run in MOPS running buffer (50 mM MOPS, 50 mM TrisBase, 3.5 mM SDS, and 0.8mM EDTA) at 150V for 50min. Gel was transferred in Biorad transfer system onto nitrocellulose membranes, blocked in 5% milk in PBS + 0.1% Triton X-100 and incubated in primary antibody overnight at 4°C. 1:10000 secondary antibody incubation was 1h at room temperature. Chemiluminescent detection with Amersham AI600, or fluorescent antibody detection with Biorad Odyssee. Western blots were quantified by densitometry using FIJI ImageJ.

### Viral infection with samples for flow cytometry, plaque assay and western blot

Cells plated the previous day were washed in PBS and infected with viral solutions in serum free DMEM at MOI of 3. At the intended time points, cell supernatants were collected and frozen at −80°C until use. For western blot samples cells were washed in PBS, immediately lysed in 2x LDS sample buffer and stored at −20°C. For flow cytometry samples cells were detached in trypsin at 37°C for 7min and transferred to a 96 well plate with conical bottom to spin down at 300xg 4°C for 3min each spin. Cells were washed once in PBS + 1% FBS and once in PBS and fixed in PBS + 4% formaldehyde for 10min at room temperature. Cells were spun down at 500xg 4°C for 5min and washed twice in PBS and stored at 4°C for a maximum of 2 days. Cells were stained for 1h in primary antibody solution or PBS for unstained controls at room temperature. Cells were spun down and washed twice in PBS and stained with 1:1000 isotype specific secondary antibody solution Alexa 488 or Alexa 647 for 30min at room temperature. Cells were spun down and washed twice in PBS and measured at Fortessa X-20.

### Flow cytometry analysis

Data acquired on Fortessa X-20 was gated for cells and single cells using FlowJo. Single cells gated data for the two measured channels M2 and HA was exported and further processed in Python with custom scripts. Optimal parameters were fitted by maximum likelihood estimation for a mixture model of two skew-normal distributions using global optimization via differential evolution. Only the distributions with the larger mean at each time point were analyzed further. The data was processed in two different ways. First, the data for each channel was normalized to the 15h time point for each protein in the WT cells and the mean at each time point was compared between WT and Δ6 cells. Second, each repeat for each protein and WT or Δ6 condition was fit to a logistic growth model using a differential evolution algorithm and normalized to the parameter L of the WT condition for each protein and repeat. The data was fit again to obtain the final optimized parameters of the normalized data.

### Plaque assay

MDCK cells were seeded in 12 well plates to reach confluency the next day. Cells were washed in PBS and infected with 300ul of serially diluted supernatants in serum free DMEM. After 1h cells were washed in acid wash solution, washed in PBS and overlayed with serum free DMEM plus 0.14% BSA and AVICEL solution. Cells were incubated at 37°C + 5% CO_2_ for 30h.

### CRISPR KO for biogenesis factors in HEK293T

Target sequences for each guide are listed in Table S3. First, two sgRNAs against cellular targets were synthesized. Gene name of human targets: NT (non-targeting), EMC3, EMC6, GET3, CAMLG, TMCO1. Genomic target sequences were designed with the chopchop web server. 2 primers were PCR amplified (25ul PCR reaction containing 0.5uM each) to build the double stranded DNA template for the in vito transcription using NEB Phusion polymerase. The forward primer contained a 5’ T7 promoter sequence (taatacgactcactatag, +G if target does not start with G), the 20 nucleotide specific genomic target sequence, and 20 nucleotides Spyro-cas9 sgRNA scaffold 3’ of the genomic target sequence. The reverse primer was the reverse final nucleotides of Spyro-cas9 sgRNA scaffold overlapping 20 nucleotides with 3’ end of the forward primer. In vitro transcription was performed with Promega T7 Ribomax Express as per manufacturers instructions in a 5ul reaction. RNA was purified on columns, eluted in RNAse free water and checked for purity and to obtain concentration on a 2% agarose gel containing 1% bleach. To perform the screen 293T cells were seeded in two wells of a 6 well plate (3×10^5 cells/well) and transfected 8h later with cas9-BLAST plasmid (1ug/well) using Jetprime reagent. 24h after transfection cells were detached with trypsin, combined and seeded in a 24 well plate (10^5 cells/well) coated with poly-D-lysine. The next morning cells were transfected with in vitro transcribed purified sgRNA (2 sgRNAs per target, each 1ug sgRNA/well in 200ul total) with Dharmafect reagent and incubated for 8h. Cells were transfected in fresh medium with M2.Y.F plasmid (300 ng/well) using Jetprime reagent and incubated for 48h. Cells were lysed in TX100 lysis buffer, separated from nuclei by centrifugation, measured for protein concentration with Bradford assay and normalized to the lowest protein concentration.

### Computational analysis of human and viral proteomes

The NCBI virus refSeq protein database (downloaded on 18.02.2025) contained 686474 protein entries. All known virus genera were obtained from ICTV (ICTV_Master_Species_List_2023_MSL39.v4) and filtered for those genera infecting humans (115 genera), by filtering for genera containing virus species known to infect humans from virus-host-db [66] (downloaded on 11.11.2024). Phages were removed from the dataset. RefSeq entries were filtered for the 115 genera to obtain 25613 protein sequences. These sequences were clustered into 10715 similar proteins with MMSeq2 [67]. Sequences were grouped together if they had at least 60% sequence identity, and covered more than 80% of an aligned sequence in the cluster (--min-seq-id 0.6 -c 0.8 --cov-mode 1). The TMBed language model [68] was used to predict transmembrane domains, their topology, and signal peptides, and transmembrane domain predictions were refined with ΔGapp calculations [10] in a 19 residue window. Transmembrane domains were considered when the location was proximal ±15 residues with a minimum in the ΔGapp calculation and was lower than 1.5 kcal/mol, resulting in a dataset of 1226 single pass membrane proteins. Protein types were manually annotated based on signal peptide presence, predicted topology, position of the transmembrane domain and length of translocated domain (typeII N-terminal length < 150 and C-terminal length > 60; type III N-terminal length < 60 and C-terminal length > 60; typeIV C-terminal length < 60; other are proteins not fitting into any of these categories). Some previously uncharacterized predicted type III proteins were validated with structure prediction.

### Statistical analysis

The statistical analysis used is detailed in each figure legend.

## ACKNOWLEDGEMENTS

This project has received funding from the European Research Council (ERC) under the European Union’s Horizon 2020 research and innovation programme (grant agreement No 101001521) to M.J.A., the BBSRC Institute Strategic Programme Grant funding BBS/E/D/20002173 and BBS/E/D20002174 to P.D. and ‘la Caixa’ Foundation project grant CF/PR/HR17/52150018 to C.A.. We thank John Christianson for the plasmids encoding SEC61B.H and SQS.H. We thank Stephen High for sharing Impomoessin F. We thank Ignacio González Bravo for critical discussions. We thank Tiago Paixão for his insights into the data analysis and guidance throughout the project.

## Author contributions

C.D., C.A., and M.J.A. conceptualized the research; C.D. and M.A. performed research; C.D., M.A., S.Z. contributed new reagents; C.D., M.A., S.Z., M.J.A analyzed data; C.D. wrote the manuscript; and C.D., C.A., P.D. and M.J.A. revised the manuscript.

