## Supplementary material for "Functional redundancy in Oxa1-family insertases supports robust membrane insertion of influenza A virus M2 viroporin": SI

SUPPLEMENTAL INFORMATION

This supplemental information file contains five supplemental figures.

Supplemental Figure S1

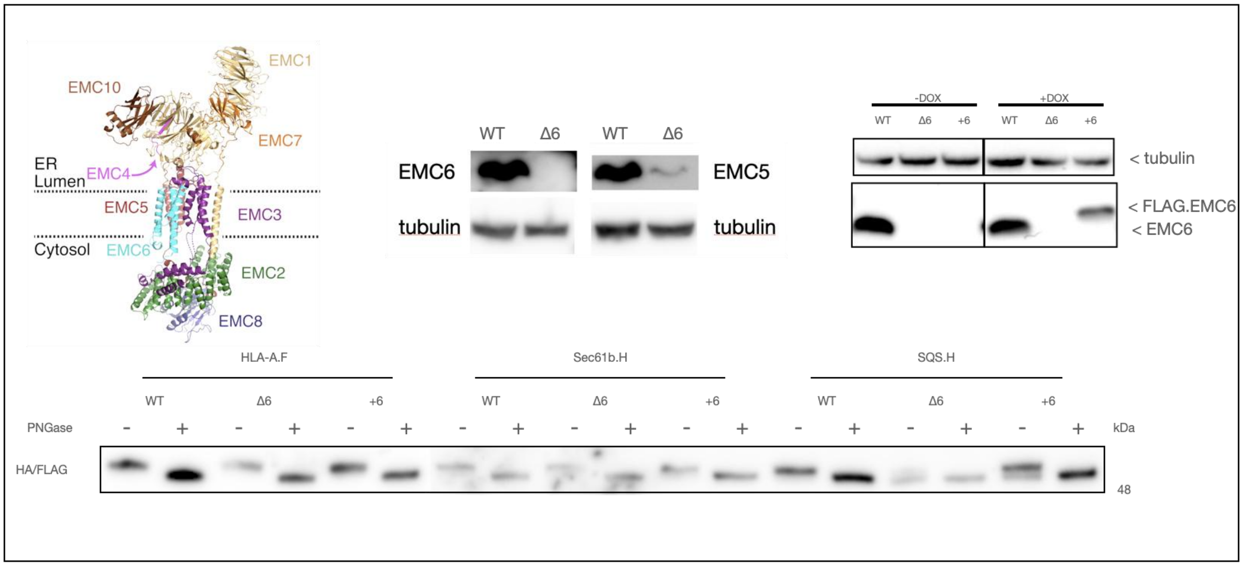


Fig. S1. A549 EMC6 KO cell line and functional validation. A549 WT, ∆6 or +6 were transfected with HLA-A.F (known Sec61 client), Sec61b.H (known GET client), or SQS.H (known EMC client) for 24. Cell lysates were analysed by western blot probed against indicated proteins. Scheme of EMC subunits is adopted from Pleiner et al. 2020.

Supplemental Figure S2

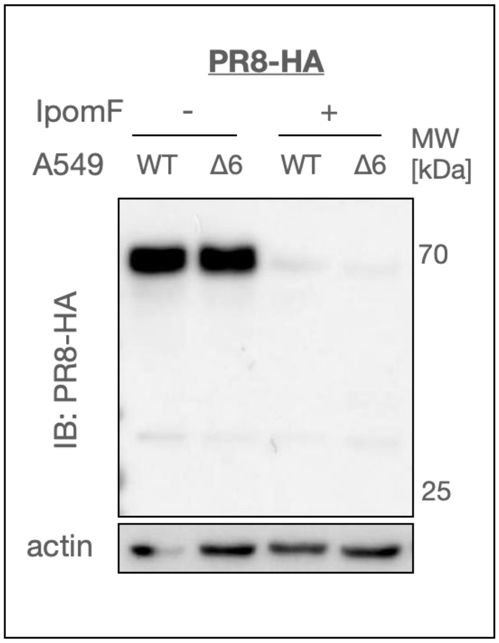


Fig. S2. IpomF validation. A549 WT or ∆6 were transfected with PR8-HA (known Sec61 client), and incubated with 50nM Ipomoeassin F (IpomF), or equivalent volume fraction of DMSO and lysates were analysed by western blot (N=1).

Supplemental Figure S2

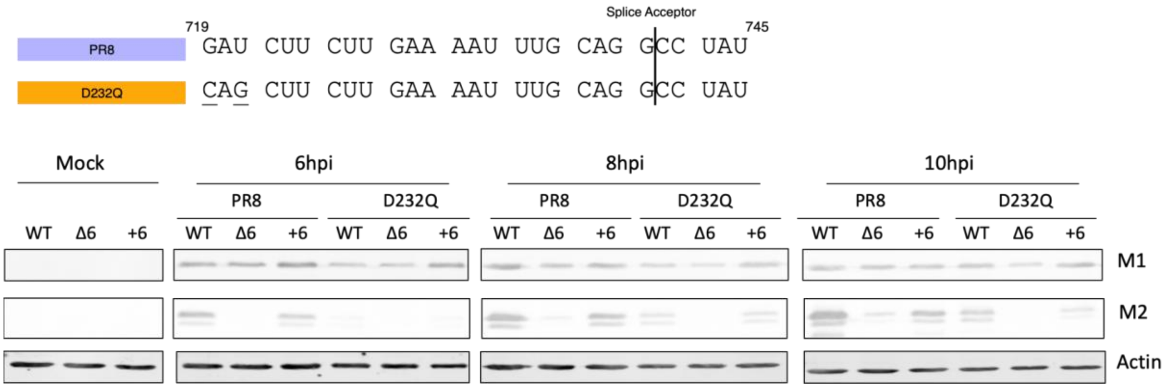


Fig. S3. EMC dependence of PR8 segment 7 splice site mutants. Scheme of the splicing adaptor sequence in segment 7. MDCK WT, ∆6, or +6 were infected with PR8 or segment 7 splice mutant D232Q at MOI 3 for the indicated times and lysates analysed by western blot (N=2).

Supplemental Figure S4

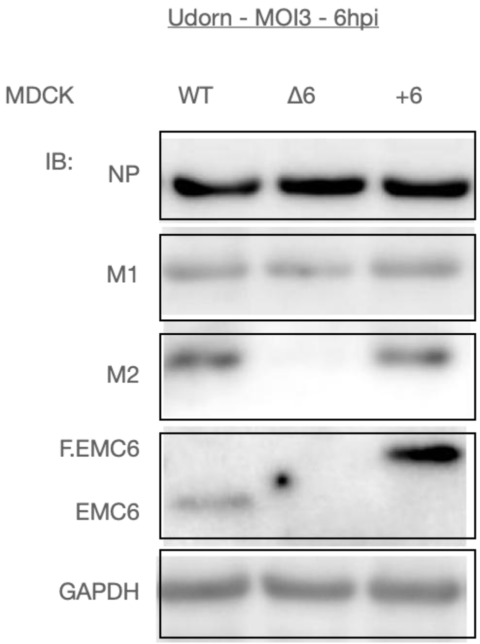


Fig. S4. Udorn viral protein expression. MDCK WT, Δ6, or +6 were infected with Udorn at MOI 3 for 6h and lysates analysed by western blot.

Supplemental Figure S5

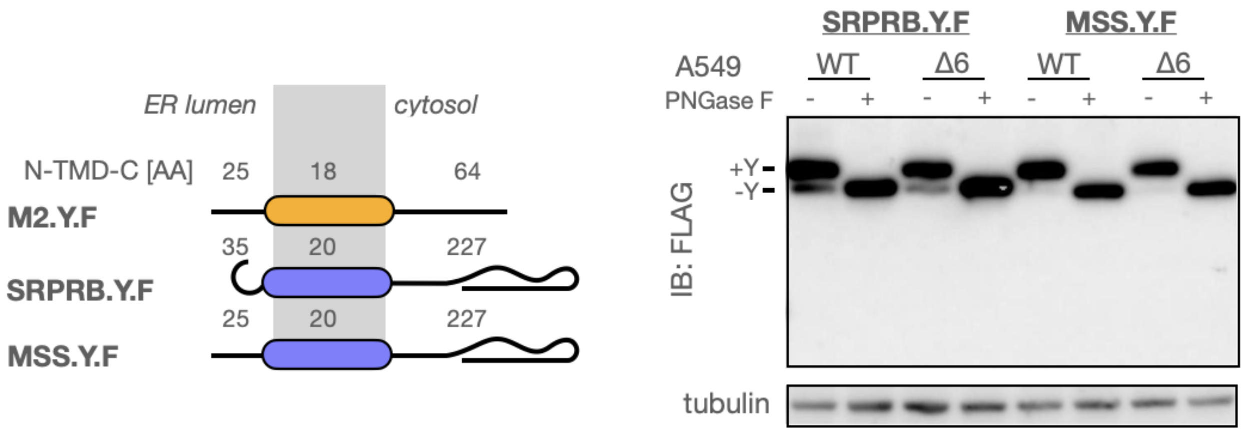


Fig. S5. M2's N-terminus does not affect EMC dependence of SRPRB. A549 WT or Δ6 cells were transfected with each of the constructs, and cell lysates were mock-treated or treated with PNGase and analysed by western blot (N=3).
